# Decoding and Characterizing the Intracranial Representation of Semantic Information

**DOI:** 10.64898/2026.07.13.738249

**Authors:** Christophe Smith, Sophie Inchyna, Bryant Barrentine, Rajesh Kana, David Basilico, Matthew Nelson

## Abstract

Brain-computer interfaces (BCIs) have achieved impressive performance by decoding motor and articulatory signals associated with speech production. However, considerably less is known about whether higher-level semantic representations can be decoded from human cortical activity. Demonstrating semantic decoding would advance both our understanding of language organization and the development of BCIs that rely on conceptual rather than purely articulatory information. We recorded intracranial neural activity from patients undergoing stereotactic electroencephalography (sEEG) for clinical epilepsy monitoring while they performed language tasks requiring semantic processing. High-gamma power was extracted from local field potentials and used to generate trial-level features for supervised machine-learning classification. Classification performance was evaluated using cross-validation. Semantic category information was decoded significantly above chance, with mean classification accuracy reaching 29.8% across 15 semantic categories (chance = 6.7%). These findings demonstrate that high-gamma activity contains information about conceptual category membership that can be extracted on individual trials. These results provide evidence that semantic information is accessible from intracranial population recordings and support the feasibility of semantic decoding as a complementary direction for future language BCIs. Beyond neuroprosthetic applications, this work contributes to understanding how conceptual knowledge is represented in the distributed human language network.

## Introduction

### Brain machine interfaces and neural decoding

A variety of conditions can result in a permanent loss of speech while leaving cognition and language skills intact. This is devastating to the individual and their support network as the effected individual must rely on those around them to manage their needs through bespoke communication tools/methods. Even if a method of communication is established, it is possible for their condition to advance, and their system of communication become unusable. This requires the individual and their support network to learn new methods of communication, but it is not uncommon for individuals to just give up and live in isolation^1^. Language-based brain machine Interfaces (BCIs) have emerged as a promising treatment for these patients as it relies only on the intact brain functions of the individual, can adapt to changes in brain function, and does not degrade to the same extent as motor function as the individual’s condition advances^2^. Recent clinical studies in patients with long term intracranial implants have shown that the system works, improves over time, and subjects are eager to use it in their daily life^3^. Most advances in this field have emerged from the use of HGP signals recorded from the motor cortex using micro-ECoG grids. These systems use low level language features such as attempted articulations decoded using machine learning models that predict attempted articulation, advanced systems pair this with a language model that predicts the most likely sequence of words^4,5^. These successes show that cortical activity contains a sufficient structure to allow accurate real-time decoding of behaviorally relevant language features.

Despite these advances, most BMI applications focus on decoding motor intentions or low-level sensory and articulatory representations. While these approaches have achieved impressive results, they primarily target execution level components of communication rather than the underlying conceptual content that guides them. As a result, the representation scope of current decoding efforts remains largely confined to the sensorimotor domain.

### Limited focus on semantics

In contrast to motor and articulatory decoding, substantially less work has focused on decoding semantic-level representations from neural activity. Here, semantic representations are defined as neural patterns reflecting conceptual content, information about meaning that is not reducible to specific sensory inputs or motor outputs. For example, recognizing a dog as a four- legged animal or retrieving the concept associated with a word involves representational information that extends beyond its visual features or articulatory form. These representations are central to cognition and communication, yet they have not been systematically characterized for use in a BMI framework.

Cognitive neuroscience has established that semantic processing engages a distributed cortical network spanning temporal, frontal, and parietal association regions^6^. Electrophysiological studies have demonstrated sensitivity to semantic categories at the population level. However, many of these studies emphasize evoked responses or group-level contrasts rather than subject-specific, trial-level decoding performance that is essential for BMI translation. As a result, it remains unclear whether semantic information can be extracted from high-frequency neural activity with sufficient reliability and generalizability to support practical decoding applications.

### Why Semantics matter

Decoding speech from the sensorimotor cortex has shown impressive performance in reconstructing attempted speech, but it targets the final execution stage of language production. Such systems do have major shortcomings. At the level of the single word decode these models struggle with selecting phonetically similar words such as fork and fort which are phonetically similar but semantically distinct. These systems perform best with continuous monitoring which patients in the target population have expressed concern about^1^, preferring more on-demand systems. Additionally, a major challenge for these systems is cued responses where subjects respond to an external stimulus such as questions, objects in the environment, cultural information, etc. This can result in ambiguous responses that a language model may struggle to understand. Semantic decoding addresses a distinct representational level, the cortical encoding of conceptual meaning. Theses representations are more distributed, temporally persistent, and less dependent on intact articulatory pathways. In individuals where the motor cortex is compromised, or for communication systems that require contextual integration rather than continuous articulatory monitoring, semantic decoding provides a complementary control dimension. Semantics is also a more robust linguistic feature abstracting away variance inherent to motor decoding such as speech rate, articulatory idiosyncrasies, coarticulation, etc. Moreover, understanding the geometry of conceptual representations in neural activity has independent scientific value for models of language organization and cognitive dysfunction. Semantic and motor decoding likely represent hierarchical components of future multi-level language BCIs. These can work in conjunction to create a BMI system that can use context and real-time attempted articulation to accurately convey speech and intended meaning in a robust system that meets real-world demands (**Figure 1**).

**Figure 1.**
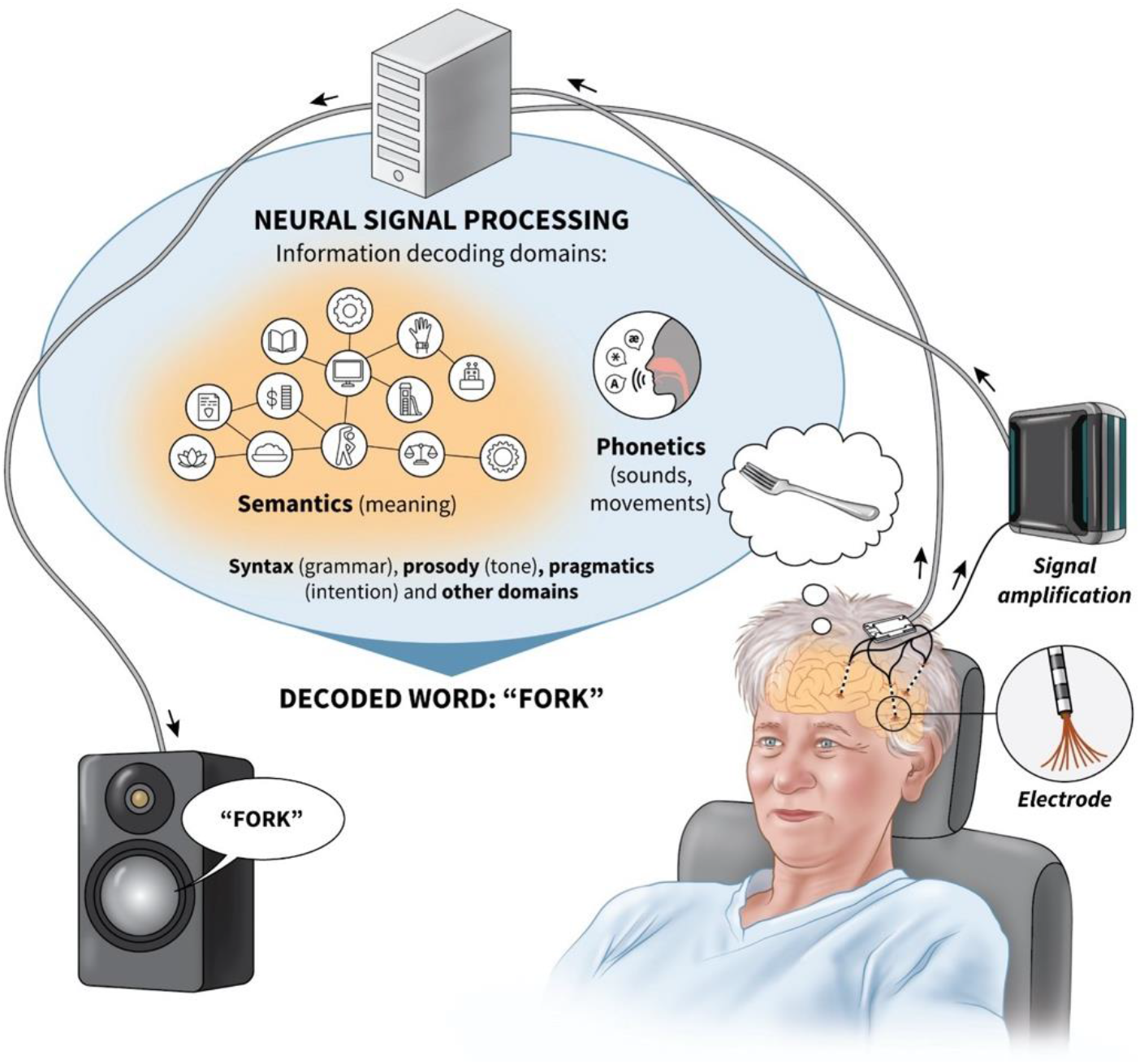
Schematic showing the potential of semantic decoding working in conjunction with phonetic and other aspects of language decoding to decode language.

### Neurological basis of semantics

Converging evidence indicates that semantic representations are supported by a distributed network across the language network. Neuroimaging, intracranial electrophysiology, and lesion studies have implicated the anterior temporal lobe (ATL)^7–10^, posterior medial temporal gyrus (pMTG), inferior temporal gyrus (ITG), posterior cingulate (PC)^11,12^, and the angular gyrus (AG)^9,13,14^ regions in semantic processing across modalities. The fusiform gyrus of the ventral ATL is a particularly crucial node in many semantic network theories^9,10,15^. The inferior frontal gyrus (IFG) is also known to play a role in semantics^6,9,12,16^ and is a strong contributor to decoding performance^17^, it is thought to play a role in executive selection of semantic information across the cortex^18–21^. Activity in these regions shows sensitivity to semantic category, conceptual similarity, and contextual meaning, suggesting they encode structured information about abstract relationships rather than purely perceptual or articulatory features. These regions are often grouped into two theoretical anatomical frameworks for the organization of semantics. The “hub-and-spoke model” proposes^22^ that the ATL is the lone central amodal hub of semantic knowledge. An alternative is what will be referred to as the “posterior model” where the posterior brain regions such as the left PC, pMTG, and ITG are amodal semantic hubs^11,23^. An alternative model is the “dual hub” model^24–28^ where the left ATL is the locus of one amodal hub specialized in taxonomic knowledge and the left AG is a second hub specialized in thematic knowledge. These models provide anatomical regions of interest (ROIs) that may encode semantic information and are promising targets for semantic decoding.

Semantics has been a long studied linguistic feature in clinical, behavioral, and intracranial research. It is commonly accepted that some brain regions specialize in processing specific semantic categories such the fusiform face area and the parahippocampal place area. Prior neuroscience studies have focused on studying the intracranial representation of a small subset of categories using fMRIs and found evidence for localized representations^22^. In clinical literature there have been documented cases of selective loss or preservation of semantic categories following focal damage to language regions^29–39^. Notably, such selective preservation of categories has not only been observed in cases of focal lesions but also in cases of extensive cortical damage, where specific categories persist despite widespread impairment^31^. In addition to this clinical evidence, there has been extensive behavioral studies characterizing semantic organization in neurologically healthy participants, demonstrating a robust and reproducible semantic organization^40^. These prior studies show causal support for structured organization within the distributed network suggesting that semantics is neither homogenous nor randomly distributed but exhibit category-sensitive organization at the cortical level. Using these studies a large selection of well characterized categories was selected that reflect well characterized categories with clinical evidence for localized representation.

High-gamma power has been widely adopted in intracranial research as a reliable proxy of task-relevant cortical activation. In many cortical regions, high-gamma amplitude correlates with local neuronal firing rates and supports accurate single-trial decoding of motor and linguistic variables. However, HGP reflects aggregate activity within a local field potential and does not directly resolve the contribution of individual neurons. Given the evidence of a distributed network with local structured populations representing semantic categories there is a risk of local population signals being obscured by more dominant distributed responses. Moreover, recent work has demonstrated that the coupling between high-gamma power and spiking activity varies across cortical regions, suggesting that HGP should not be assumed to uniformly capture underlying neuronal selectivity^41^.

### Methodological opportunities

Electrocorticography (ECoG) and stereotactic EEG (sEEG) recordings offer millisecond temporal precision and substantially higher signal-to-noise ratios than non-invasive techniques enabling reliable estimations of task-related neural dynamics at the level of cortical populations. HGP (70-150Hz) has been shown to correlate with local neuronal firing rates providing a physiologically interpretable feature space for decoding.

Much of the major advancements in motor-based language BCI systems has been done using chronically implanted micro-ECoG arrays^4,42–44^. However, the use of such grid arrays has fallen out of favor in epilepsy centers in the United States being replaced by sEEG depth electrodes. The patient population that qualifies for the micro-ECoG arrays used in prior studies is far smaller than the epilepsy population. This has created a need for studying new BCI development platforms that match common clinical practices and works with a large and readily available clinical population.

Developments in computational methods now allow systematic extraction and evaluation of high-dimensional neural features. Time-resolved feature extraction pipelines permit quantification of HGP activities across electrodes and trials, while machine learning classifiers enable testing if these features contain discriminative information about experimental conditions. Vitally, cross-validation frameworks and feature selection procedures reduce overfitting and allow assessment of model generalization on hold-out data. These tools transform decoding from a descriptive analysis into a statistically principled inference problem.

These advances make semantic-level decoding possible. By employing controlled stimuli while collecting neural data and implementing a cross-validated classification framework with feature selection. The present study directly tests whether HGP encodes semantic features rather than low-level sensory or task related confounds. This bridges theoretical models of semantics with rigorous empirical implementation.

Despite extensive characterization of the cortical networks supporting semantic processing and substantial progress in motor and speech decoding, it remains unclear whether semantic-level representations can be reliably decoded from high resolution intracranial activity in a manner that supports generalizable BMI relevant classification. It is unknown if conceptual information can be distinguished at the single-trial level independent of motor and task confounds, and whether such decoding generalizes across example objects within a semantic category. It is also unknown if this semantic representation has a consistent amodal cortical representation.

In this study, we investigate whether semantic category information can be decoded from intracranial neural recordings using high-frequency population measures of activity to generate features. The stimuli used is a set of concrete nouns that utilize multiple modalities and establish the feasibility of semantic decoding. Participants engaged in tasks designed to elicit semantic processing and isolate the comprehension signal from motor responses. We extracted HGP and spike frequency derived features from the distributed regions in the language network and used supervised machine learning models to classify semantic categories at the single-trial level. Critically, we evaluated generalization performance across stimuli to determine whether decoding reflected category-level structure rather than item-specific features.

We hypothesized that high-frequency activity within the language network would contain sufficient information to enable above-chance classification of semantic categories, and this decoding could generalize across items in the same category. Demonstrating reliable decoding would provide evidence that conceptual representations can be operationalized within a BMI framework and may offer a complementary tool to motor-based approaches.

To our knowledge, few studies have directly evaluated semantic category decoding from intracranial recordings using this approach. The outcomes of this study show that this type of language decoding is safe and feasible laying the groundwork for studies with a broader scope and longer implantation periods. It will also address several issues of scope in semantic decoding such if there are significant differences across modalities and the granularity of semantic concepts in the cortex.

## Methods

### 3.1 Ethics and oversight

All protocols involving human participants were approved by the Institutional Review Board (IRB) at the University of Alabama at Birmingham (Protocol #300004963). Written informed consent to do the tasks was obtained from all participants prior to enrollment in the study. All procedures were conducted in accordance with the Declaration of Helsinki.

Data was collected as part of a registered clinical trial (ClinicalTrials.gov Identifier: NCT05283265). The present analysis constitutes a secondary computational analysis and is not designed to evaluate clinical endpoints.

### 3.2 Experimental Design

#### 3.2.1 Patient population

Over the course of the study participants were recruited from patients undergoing clinical care at the University of Alabama at Birmingham Heersink School of Medicine Epilepsy Monitoring Unit (EMU) from March 2023 to February 2026. Inclusion criteria required that patients (i) had a confirmed diagnosis of pharmacoresistant focal epilepsy requiring long-term intracranial monitoring, (ii) were ≥ 18 years of age at the time of recording (iii), had no prior resections or major structural brain pathologies, (iv) were mono-lingual native English speakers with nominal language and cognitive skills. Electrode implantation was chosen by the patient’s clinical care team to meet clinical demands; trajectories were planned to maximize coverage of the language when feasible. Electrode location was not used as in inclusion criterion for patients, but electrodes were excluded based on ROI and if the task is unresponsive to the task. Participant data was excluded if (i) they failed to complete 1 of the 2 research sessions or (ii) data collected during the trial was rendered unusable due to a technical fault of the task or recording equipment. Patients remained in the EMU for 3 – 28 days completing both sessions over a 2-to-7-day period depending on patient availability.

#### 3.2.2 Electrode Design and Data Acquisition Electrodes

Depth electrode recordings were obtained using sEEG electrodes from Ad-Tech Medical Instrument Corporation. Each electrode consists of a polyurethane shaft with 8-16 platinum-iridium contacts embedded in the surface (contact length 2.0mm; inter-contact spacing 1.5mm). These electrodes are placed primarily for clinical monitoring of pharmacoresistant epilepsy, the electrode targets and trajectories were solely determined by clinical considerations. Each channel is re-referenced to a hardware reference recorded from a dedicated reference electrode.

Subdural ECoG recordings were obtained using platinum grid and strip electrodes from Ad-tech Medical Instrument Corporation. Grid electrodes consisted of a silicone substrate with 4-36 circular platinum-iridium contacts (diameter 4mm; exposed surface 2.3mm; center to center distance of 5mm) embedded in the substrate arranged in a variety of strip and array configurations. Electrodes were placed on the cortical surface as part of clinical monitoring with placement determined by clinical considerations. Each channel is re-referenced to a hardware reference recorded from a dedicated reference electrode.

Data acquisition platforms:

sEEG and ECoG, were recorded using a clinical EEG acquisition system (Natus Quantum, Natus Medical Incorporated) and exported in EDF format. The amplifier has an input impedance of >1 GΩ and signals were sampled at 2,048 Hz with a 24-bit resolution. The exported EDF files reflect the raw recordings and include the acquisition systems front-end acquisition filtering (low-frequency high-pass and anti-aliasing low-pass filters) applied prior to digitization. Clinical display band-pass filters and montages were not included in the exported data.

All data was acquired in the patients EMU room.

#### 3.2.3 Behavioral Testing Software

Behavioral tasks were implemented using E-Prime (version 3.X; Psychology Software Tools) and administered on a touchscreen laptop computer (1920x1080). Visual stimuli were presented on the integrated display (60 Hz refresh rate) and audio stimuli was presented with the integrated speakers. Responses were recorded as touch responses using the capacitive touch screen or audio responses recorded on two external microphones. One microphone is connected to the computer and records audio through the E-Prime software at a sample rate of 80 kHz. The second microphone acted as a back-up and synchronization tool and was connected to an amplifier and then a direct current channel (DC) on NATUS with a sample rate of 2048 Hz. All patient rooms were under 24/7 audio-visual monitoring recordings made during session was collected to act as a final fall back if data was lost. In tasks that are paced by the experimenter a Bluetooth clicker was used to advance the slide after the patient responds. Over the course of the study, multiple touchscreen laptop models were used; however, all systems provided comparable display sizes and resolutions, and identical task scripts and timing parameters were used across devices. Over the course of the study the operating system used was upgraded from windows 10 to windows 11.

The behavioral data was synchronized with the neural data acquisition systems using a signal box (BrainTrends). The signal box was modulated by E-Prime sending 5 V TTL pulses using in-line scripts and E-Prime events. The signal was recorded in NATUS using a DC channel. If the ATLAS system was used the signal box’s output was split and connected to a NATUS and an ATLAS DC channel.

### 3.3 Data Acquisition and Signal Processing LFP processing

Local field potentials were processed offline for macro-contact signals, these signals re- referenced using a bipolar montage^16,45^. Bipolar pairs were determined using electrode nearest neighbors excluding channels that were removed due to channel artifacts. The location of these new bipolar signals is treated as being in the midpoint location between the electrode pair in MNI coordinates. Going forward, all references to electrodes refer to these re-referenced signals. Signals were then processed using wavelet spectral analysis (Field Trip toolbox in MATLAB^46^) to compute signal power across the frequency range the wavelet bias is corrected by multiplying the transform by the frequency. Then for all bins used in the analysis the power is averaged.

Finally, the signal is then scaled based on baseline activity using a z-score of trial data relative to the entire session. This method is used to calculate the instantaneous high-gamma power (HGP) using frequencies from 70-150 Hz and 26 lower frequency bands spaced logarithmically from 2- 70Hz^47–49^.

### 3.4 Task Design

#### 3.4.1 Method: Semantic category selection

Semantic categories were selected a priori based on three criteria. First, categories needed to have evidence for localized representations in clinical literature and needed to be thoroughly characterized in behavioral studies. Second, categories were restricted to concrete concepts that have members that are unambiguous across text, audio, and visual modalities. Third, categories were chosen so that semantic categories are the lowest level of a nested semantic hierarchy that matched the broad categories studied previously. The 15 categories chosen are four-footed animals, bugs, human professions, human body-parts, man-made foods, fruits or vegetables, musical instruments, vehicles, kitchen ware, articles of clothing, famous people, buildings, colors, and shapes. These categories can be organized into the following superordinate categories artifactual objects (musical instruments, vehicles, tools, kitchenware, articles of clothing), people (famous people, human body-parts, human professions, articles of clothing), places (buildings, named buildings), faces (famous people), low level concepts (colors, shapes), animate (four-footed animals, bugs, human professions, body parts), and inanimate (man-made food, fruits and veg, tools, musical instruments, vehicles, kitchen ware, articles of clothing). This results in 15 semantic categories, 7 superordinate categories, and 3 levels of semantic organization to study, the main hierarchy used in this study is shown in **Table 1**.

**Table 1.** Stimulus categories and their associated hierarchy.

| **Hierarchy of Semantic Categories** |
|---|
| Animate |
| Four-Footed Animals (cat, cow, etc.) |
| Bugs (spider, ant, etc.) |
| Human Professions (plumber, teacher, etc.) |
| Famous People (Einstein, Gandhi, etc.) |
| Human-Body Parts (arm, ear, etc.) |
| Inanimate |
| Artifacts |
| Musical Instruments (piano, violin, etc.) |
| Vehicles (airplane, car, etc.) |
| Tools (hammer, screw, etc.) |
| Kitchenware (plate, fork, etc.) |
| Articles of Clothing (shoe, sweater, etc.) |
| Man-Made Foods (taco, pizza, etc.) |
| Fruits and Vegetables (apple, pumpkin, etc.) |
| Places |
| Buildings (house, church, etc.) |
| Low level concepts |
| Colors (blue, red, etc.) |
| Shapes (square, circle, etc.) |

#### 3.4.2 Method: Object selection

Each category had 15 representative objects that met the following 4 criteria. First, the object must be recognizable as an image. Second, the object should be unambiguous in all modalities. Third, where possible noun-noun polysemia was avoided, this rule does not apply to noun-verb polysemia since the task is restricted to nouns (E.g. saw). Finally, unless the objects name is a proper noun the object must have a single word representation, compound nouns were allowed (E.g. stick bug would not be allowed, but ladybug is allowed). For some categories exceptions were made due to the limited number of recognizable category members. If an exception was made the object would only be used for one category. Images of common noun objects were presented as colored illustrations proper nouns were presented as photos or portraits. in the text modality all objects are presented as lower case text that occupies a single line on our testing monitor, and in the audio modality a recording of each word was made by a native English-speaking human male with clear enunciation and no background noise. All audio was presented for a minimum of 1 second, longer audio was allowed to play until it finished.

Two stimuli sets were used in this task where each set controlled for different linguistic features. The first set was designed to use canonical category objects; these are objects that our research population are likely to think of as being a highly representative member of a given category. These objects were selected from stimuli lists developed in prior studies and by internal lab discussion. This set was chosen to maximize neural response to objects that clearly represent a category and statistical imbalance of the data set would be addressed post-hoc. The second set was designed to balance psycholinguistic linguistic features pre-hoc. The features considered for common nouns were word frequency^50,51^, word length, syllable count^52^, phoneme count^51^, orthographic neighborhood size^51^, and bi-phone frequency^53^. Each psycholinguistic feature for common nouns was balanced so that the ANOVA/Kruskal-Wallis p value was > 0.1. This resulted in 2 stimuli sets with 225 objects across 15 categories.

### 3.5 Task overview (production)

Three production tasks were used in this study (**Figure 2**). These tasks require participants to translate ideas into language with each task requiring participants to perform some form of categorization.

**Figure 2.**
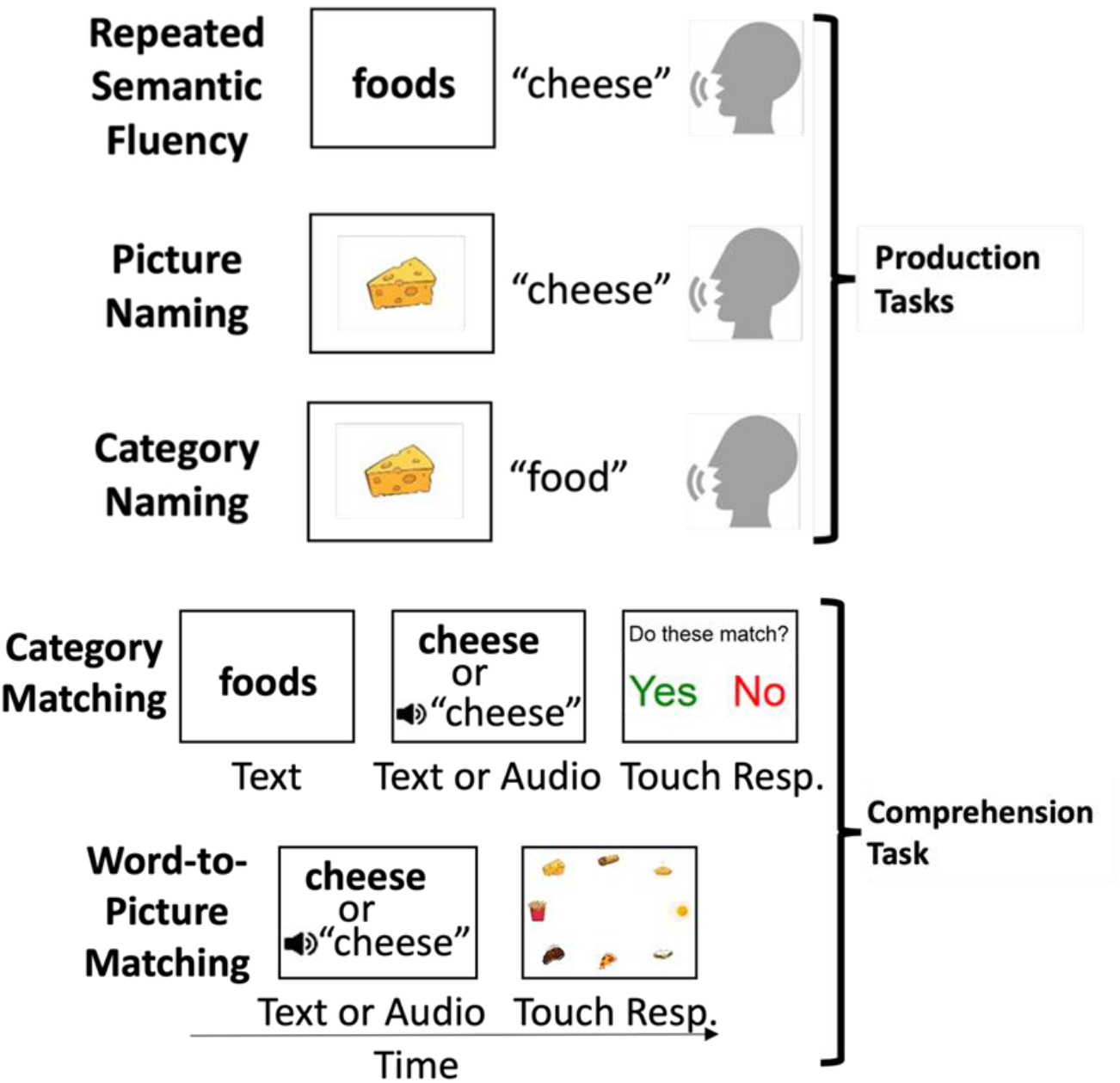
Schematic showing the 5 tasks studied.

#### 3.5.1 Picture Naming task (PNT)

During the Picture Naming Task (PNT) participants are shown an image of an object at the center of the screen. Objects that are common nouns are shown as colored illustrations in the center of the screen. Proper nouns are shown as photos or portraits that have the focus in the center of the screen, and the image takes up the full screen. Participants are tasked with saying the name of the object out loud as fast as possible while their response is recorded by the testing computer. There are 225 trials of this task, and it is split into 5 blocks with 45 trials par block.

#### 3.5.2 Semantic Category Naming task (SCNT)

The Semantic Category Naming Task (SCNT) proceeds in the same way as PNT, but participants are asked to state the category they believe the object belongs to. There are 225 trials of this task, and it is split into 5 blocks with 45 trials par block.

#### 3.5.3 Repeated Semantic Fluency Task (RSF) During Repeated

Semantic Fluency Task (RSF) a random category is presented as lower case text at the center of the screen. The participant is then tasked with giving an example item from within that semantic category. If the item is new in the current session the experimenter will advance the task to the next trial. If the item is not new the experimenter will ask them to come up with a new unused item in the category. The participant is allowed to keep responding until they come up with a new item or they give up. If the participant comes up with a new item the experimenter will advance the task to the next slide if they cannot come up with a new item, then the semantic category is removed by the experimenter. The stimuli list contains the 15 categories locally balanced with no back-to-back repeats the list is composed of 840 trials. The task is time controlled and participants do 5 blocks of 5 minutes completing an average of 200 trials over the whole session. The stimuli list is pre-generated so as categories are removed there is a risk of back-to-back repeats. If all 15 categories are removed, then the task will not reoccur in the current session. This task probes participants internal representations so they are given a level of leniency with their responses. The experimenter will not try to prompt different responses even if they disagree with the categorization.

This task is modeled after the traditional semantic fluency task used by neuropsychologists but altered to run in a trial-by-trial format. This change allows for repetition over many trials, and it allows for the dynamics of neural representation to be studied as each trial requires the participant to determine the target category and search for a new member of the category.

The response is recorded by the testing computer that records all responses for each trial. The experimenter will record responses on a second computer to track the participants response and find repeated responses. Offline the audio is labeled by lab members who use the experimenter’s notes and the audio to determine the participants responses. Leeway is given to the participant since we are probing their internal organization. There are cases where users will be obviously incorrect such as cases of semantic paraphasia or neologisms.

### 3.5 Task overview (comprehension)

Two comprehension tasks were used in this study. These tasks require participants to translate language into meaning with each task requiring participants to identify some form of conceptual matches.

#### 3.5.4 Word-to-picture task (W2P)

During the Word-to-Picture task (W2P) participants are given a cue word as lower-case text or audio played by the test computer. This is followed by a slide with a search array with 8 items arranged in an ellipse around the center of the screen. One item matches the object in the cue screen the remaining seven items are distractors chosen from the same category as the cue item. Responses are recorded by having the participant touch the target image in the search array and the test computer records their response. The stimuli are balanced so that all items are present in the search array 8 times for the text modality and 8 times for the audio modality with no images being duplicated in a single search array. The modality swaps every 37-38 trials, and each block has 90 trials, so within a block the modality swaps 3 times.

#### 3.5.5 Semantic category matching task (SCM)

During Semantic Category Matching Task (SCM) participants are presented with one of the 15 semantic categories as lower case text at the center of the screen for 650ms. Then an object is presented as either lower-case text in the middle of the screen or audio played through the testing computers built in speakers. Images are on screen for 1 second and the audio slide is on screen for a minimum of 1 second but is long enough to play the full audio. The subject is then given a question slide asking if the object is a member of the presented category. Their response is recorded by using the test computers touch screen to touch “yes” or “no” on the monitor. Every object in our stimuli set is provided as text and audio so there are 225 audio trials and 225 text trials resulting in a total of 450 trials for one session of SCM. For half of the trials the object will belong to an item in that semantic category, then for the remaining half it will be outside of that category as either a distant error, semantic categories are not similar, or a similar error where the object category and the shown category are related but distinct.

#### 3.5.6 Task Structure

Participants completed the 5 tasks over the course of 2 sessions lasting 1-1.5 hours. Each task probes different components of semantic representation but they have similar components that are interesting to use in direct comparisons. To facilitate this direct comparison the tasks were formatted in a manner designed to normalize variations in subject arousal and attention.

Each day had 2 production tasks (one is always RSF) and 1 comprehension task. Each task was split into smaller blocks with 20% of the task’s stimuli set in each block. Subjects would do a block of each task starting with the first production task, then the comprehension task, then the second production task. After completing a pass of all three tasks they loop back to the start and do the same loop again with the next block of tasks. The target time for each block is 5 minutes, each cycle of 3 task blocks takes around 15 minutes, and finishing all 5 blocks of each task takes 1.25 hours. Between subjects the specific task order can vary across 4 preset task orders explained in S2**.3**. Between cycles participants would be prompted to take short breaks with one required break after cycle 3.

#### 3.5.7 Behavioral data preprocessing

During all session notes are taken that track information about responses, distractions, and any factors that may impact trial viability. In addition, audio-visual data collected from the room is used to review the session and resolve issues in the notes or the behavioral data. If a trial is deemed unusable due to an issue during the experiment it is excluded from all future analysis. After removing trials that have an experimental flaw all audio is processed and labeled. This is done by using a local machine learning model (MATLAB’s wav2vec) to make initial predictions and onset detections. The transcriptions are then checked by a human reviewer who goes through and adjusts the labels and onset to match the data. A random sample of the labels are then reviewed by another lab member who note revisions and mistakes and sends it back to be relabeled. This continues until the labels pass inspection at which point the labels are stripped from the audio data and used for analysis. These labels include qualitative information about the trial alongside the actual start time, stop time, and transcription.

### 3.6 Computational Analysis

#### 3.6.1 Signal Preprocessing

To reduce signal noise and normalize data across trials each trials several preprocessing steps. To reduce signal noise, a gaussian convolution was applied to each channel with a standard deviation of 0.075s. To correct for global drift a baseline correction was used by taking the average power in a 0.6s baseline window measured before the start of the trial and subtracting that from the task signal.

#### 3.6.2 Feature Extraction

For a given trial an analysis window was defined based on a start event that all trials were aligned on and a final event. The analysis window was defined as starting 200ms after the start event and ending 50ms after the stop event. To ensure all analysis windows had the same time scale and produced the same number of features all windows were truncated to match the length of the shortest trial included in analysis. The truncation was done on the end event edge where temporal alignment is worst. Within this analysis features were extracted from bins tiled across the analysis window with a bin size of 200ms and a step size of 50ms. The data within these windows was then averaged to get an average HGP for the given window. The set of features derived from these windows was treated as an independent feature vector during classification. Features were normalized by z-scoring the features used in the training set and the training set was z-scored using the mean and standard deviation of the test set to ensure consistent feature meaning across training and testing.

For each trial and time window the feature vector consisted of derived values from all electrodes in each patient unless otherwise specified. To select high variance features the training set was analyzed using ANOVA to select the features with a p-value < 0.1, no testing features are used in this process to avoid letting testing information leak into the training set. The selected feature indices were used to select features from the testing set so that the features used to train and test the model are consistent. Following selecting features, the features for the entire trial were merged into one representative vector and used to train the model. The test set was then used to evaluate the model.

#### 3.6.3 Data Partitioning and Cross-Validation

Classification was performed within subject within a specific task. Trials were partitioned using stratified five-fold cross validation. In each fold, 80% of the trial was used for training and hyper parameter optimization, and 20% was held out for evaluation. Within training sets data was up-sampled so that each category had equal representation, but up-sampling was not applied to the test set.

Hyperparameter tuning was performed using nested five-fold cross-validation within the training set of each outer fold. Parameter selection was based on the median balanced accuracy across inner folds.

To prevent data leakage all preprocessing steps that could introduce data leaks (normalization, augmentation, hyper parameter selection) was confined to the training data of each fold. Test data was not used to make determinations about training data, rather where appropriate parameters selected using the training data was applied to the test data to ensure features represent the same information across training and testing.

#### 3.6.4 Model Specifications

A support vector machine (SVM) with a linear kernel was used to analyze the derived feature set. A linear kernel was chosen due to the high feature dimensionality relative to sample size, the interpretability of the weights, and its robustness against over fitting. L2 regularization was applied with the soft max parameter C (0.1).

#### 3.6.5 Hyperparameter optimization

For hyperparameter optimization parameters were searched using a grid search within nested-cross validations. Parameters ranges searched were: SVM regularization parameter C: {10^-3, 10^-2, 10^-1, 1, 10, 10^2, 10^3}, temporal smoothing windows 50-150ms with 50ms step sizes, and baseline correction. Within each fold, hyperparameters were selected based on the median balanced accuracy across the inner folds. The best performing hyperparameter combination was then used to fit a model to the entire outer training set, and the full model was evaluated on the held-out test set. An exhaustive list of search parameters and their results are given in S5.

#### 3.6.6 Classification Procedure

The SVM was trained and minimized loss using hinge loss with L2 regularization. Decisions were made using a one-vs-one SVM consensus mechanism.

#### 3.6.7 Preregistration/ Model development

To mitigate model bias, model development was restricted to a pre-selected subset of subjects (development cohort). Within this cohort, model architecture, parameters, and classifier hyperparameters were iteratively optimized using cross-validated performance as the optimization metric. Once an optimal configuration was found, all preprocessing and hyperparameter choices were fixed. These fixed specifications were then applied without modification to an independent set of held-out subjects (evaluation cohort). For this cohort training, hyperparameter optimization, and performance were conducted following the identical cross-validation procedure. No evaluation data was used during the development phase.

### 3.7 Statistics

Unless otherwise noted, the decoding performance is reported as median ± the inter quartile range (IQR) across results. The statistical significance of label classification relative to chance was assessed using permutation tests (1000 label shuffles per task). Empirical p-values were calculated as the proportion of permutations yielding performance equal to or greater than the observed value.

Comparisons between conditions were performed using two -sided paired t-tests when normality assumptions were met.

To ensure the model identifies class-specific patterns and is not mapping high- dimensional noise to the categories a false discovery rate (FDR) correction is applied using Benjamini-Hochberg correction^54^. The category decoding performance was deemed significant at an FDR of 5% (q<0.05)

All statistical analysis were performed in MATLAB R2024b using the statistical toolbox.

## Results

Example responses from individual recording sites are shown in **Figure 3**. These representative channels illustrate that individual recording locations exhibited distinct response profiles across semantic categories rather than uniform activation. For example, one electrode located in the posterior temporal cortex showed relatively strong responses to categories including famous people, articles of clothing, and vehicles, whereas a second recording site demonstrated greater responses to kitchenware and fruits and vegetables. These examples illustrate that category selectivity can be observed at the level of individual recording sites while also highlighting the heterogeneous organization of semantic representations across the cortex.

**Figure 3.**
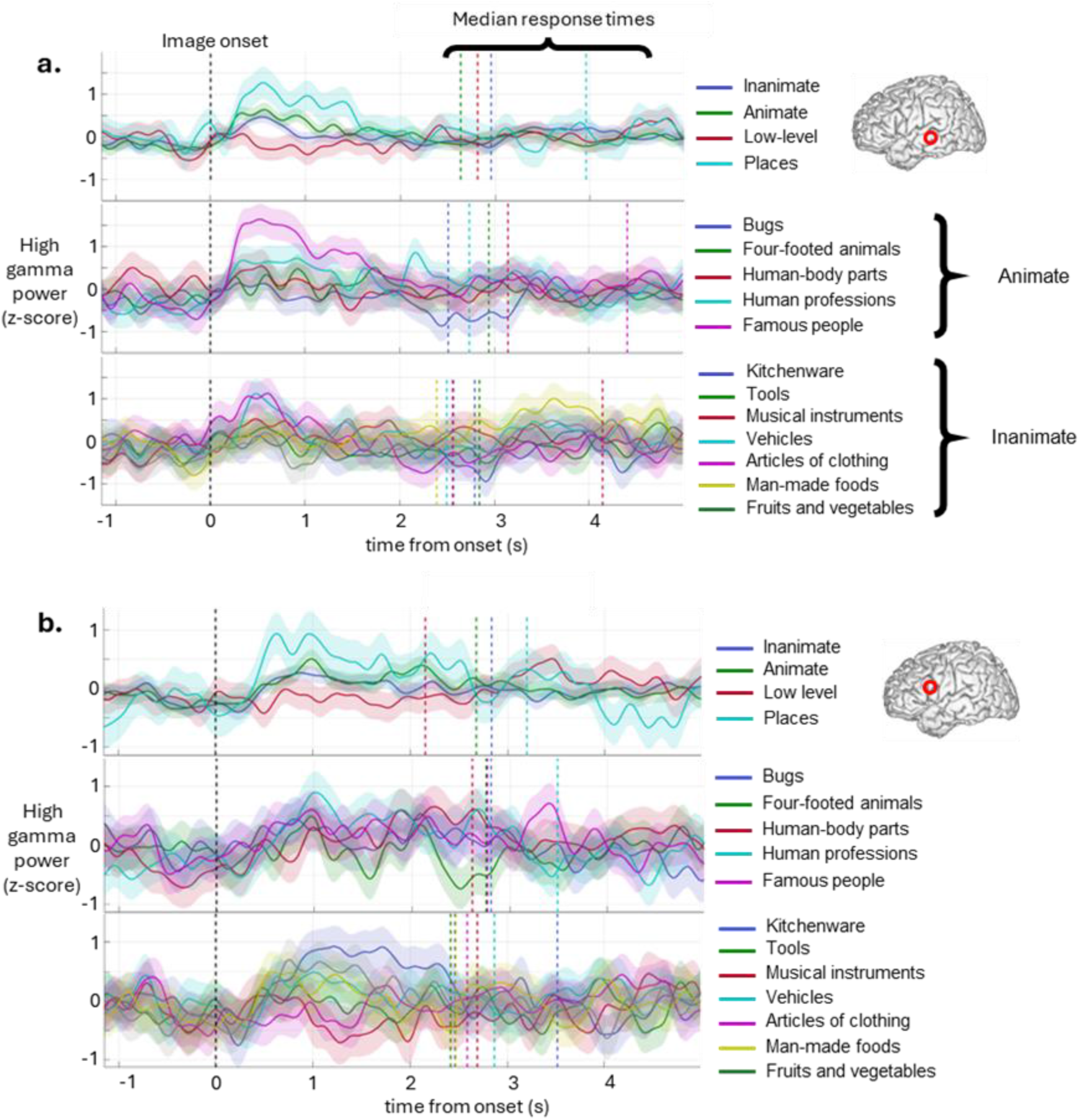
Single channel neural activity recorded in two locations from the same patient. (A) High levels of discrimination shown between animate semantic categories. (B) High levels of discrimination shown between inanimate categories. Both locations show separation between superordinate categories.

To determine whether these category-dependent responses could be used for decoding, we trained supervised machine-learning classifiers using high-gamma power features extracted from intracranial recordings. Classification was performed independently within subjects using cross-validation procedures designed to minimize overfitting.

To evaluate decoding performance across semantic language tasks, classification models were independently optimized for each participant and task using nested cross-validation.

Hyperparameter optimization included the SVM regularization parameter (C) and the ANOVA feature-selection threshold. The optimal parameter combination for each participant-task combination was then used to evaluate classification performance. The resulting decoding accuracies and selected model parameters are summarized in **Table 2**.

**Table 2.** Decoding accuracies for each patient-task combination. Blank cells indicate where data was unavailable for the corresponding patient–task combination.

| **Repeated Semantic Fluency** |  | **Category Naming** |  | **Picture Naming** |  | **Category Matching** |  | **Word-to-Picture** |  |
|---|---|---|---|---|---|---|---|---|---|
| Patient | Decoding accuracy (%) | Patient | Decoding accuracy (%) | Patient | Decoding accuracy (%) | Patient | Decoding accuracy (%) | Patient | Decoding accuracy (%) |
|  |  | P1 | 32.57 | P1 | 22.48 | P1 | 36.38 | P1 | 32.48 |
| P2 | 32.18 | P2 | 27.11 | P2 | 28.71 | P2 | 20.31 | P2 | 26.67 |
| P3 | 11.35 | P3 | 22.13 | P3 | 11.73 | P3 | 14.16 | P3 | 10.27 |
| P4 | 53.46 | P4 | 22.49 |  |  | P4 | 27.01 |  |  |
| Mean: | 32.33 | Mean: | 26.08 | Mean: | 20.97 | Mean: | 24.47 | Mean: | 23.14 |

***Chance level: 6.67 %***

Across the evaluated tasks, decoding performance consistently exceeded the chance level of 6.7%, although performance varied across both participants and task paradigms. The highest decoding accuracies were observed during the Repeated Semantic Fluency task for Participant 4 (53.46%) and the Category Matching Task for Participant 1 (36.38%).

Together, the single-channel examples and population decoding analyses indicate that semantic category information is represented in distributed intracranial neural activity at a level that supports reliable machine-learning classification. These results establish the feasibility of decoding conceptual semantic information from intracranial recordings using high-gamma features.

## Discussion

In this study, we investigated whether semantic category information can be decoded from intracranial recordings of human cortical activity. Using high-gamma power derived from local field potentials recorded during multiple semantic language tasks, we found that semantic category membership could be classified substantially above chance. These findings demonstrate that conceptual information is represented in population-level cortical activity at a level sufficient to support single-trial machine-learning classification.

Most previous language BCIs have focused on decoding articulatory movements or low- level speech representations from sensorimotor cortex. Those approaches have achieved remarkable performance, but they primarily target the motor execution stage of language production. In contrast, semantic decoding targets conceptual representations that precede articulation and may therefore provide complementary information for future language BCIs.

Beyond potential neuroprosthetic applications, the ability to decode semantic information also provides a new experimental framework for studying how conceptual knowledge is represented across distributed cortical networks.

Several limitations should be considered. The present study evaluated decoding performance in patients undergoing temporary intracranial monitoring for epilepsy, and recordings were limited by clinically determined electrode placement and recording duration. Furthermore, the analyses focused primarily on high-gamma power derived from local field potentials. Future work incorporating single-neuron activity, additional neural features, larger participant cohorts, and more sophisticated decoding approaches may substantially improve performance and provide greater insight into the neural organization of semantic representations.

Overall, these findings demonstrate the feasibility of semantic decoding from intracranial recordings and establish a foundation for future work investigating conceptual representations in the human brain. Future studies will examine additional semantic relationships, compare semantic representations across sensory modalities, evaluate the contribution of individual cortical regions, and explore how semantic decoding can be integrated with existing motor-based language BCIs to improve communication neuroprostheses.

## Acknowledgements

This work was supported by NIH grants K01DC019165 and K01DC019165-03S1 (MJN). CS was supported by NIH institutional training grant T32NS061788.

